# Quiescence unveils a novel mutational force in fission yeast

**DOI:** 10.1101/126755

**Authors:** Serge Gangloff, Guillaume Achaz, Adrien Villain, Samia Miled, Claire Denis, Benoit Arcangioli

## Abstract

**One Sentence Summary:** The quiescence-driven mutational landscape reveals a novel evolutionary force.

**Abstract:** During cell division, the spontaneous mutation rate is expressed as the probability of mutations *per* generation, whereas during quiescence it will be expressed *per* unit of time. In this study, we report that during quiescence, the unicellular haploid fission yeast accumulates mutations as a linear function of time. We determined that 3 days of quiescence generate a number of invalidating mutations equivalent to that of one round of DNA replication. The novel mutational landscape of quiescence is characterized by insertion/deletion accumulating as fast as single nucleotide variants, and elevated amounts of deletions. When we extended the study to 3 months of quiescence, we confirmed the replication-independent mutational spectrum at the whole-genome level of a clonally aged population and uncovered phenotypic variations that subject the cells to natural selection. Thus, our results support the idea that genomes continuously evolve under two alternating phases that will impact on their size and composition.

## Introduction

The causes and consequences of spontaneous mutations have been extensively explored. The major sources include errors of DNA replication and DNA repair and the foremost consequences are genetic variations within a cell population that can lead to heritable diseases and drive evolution. The knowledge of the rate and spectrum of spontaneous mutations is also very informative and of fundamental importance to understand their origin. During cell division, fluctuation assay (1-2) and more direct measurements using next generation sequencing, including mutation accumulation (MA) (3) and *de novo* mutations (4) improved the mutation rate estimations expressed as the number of spontaneous mutations *per* generation and led to the definition of their spectrum in many species (5-7). During growth, most of the mutations are due to DNA replication errors. When the mutation is neutral or beneficial, it is fixed in the population during the following round of DNA replication. Conversely, when the mutation is disadvantageous it is rapidly counter selected. The replication-dependent mutations define a spectrum quite similar among species, with the domination of Single Nucleotide Variants (SNVs) over Insertions/Deletions (Indels) and Structural Variants (SVs). For instance, preference for insertions at the expense of deletions along with a universal substitution bias toward AT has been frequently reported during cell division (8, 9). At the evolutionary timescale, the mutations accumulate as a linear function of time (10). However, the mutation rate is not constant and depends on the generation time, the efficiency of the DNA damage protection, the accuracy of DNA repair and the environment. In the past decades, it has become increasingly evident that mutations also arise during cell cycle arrest, slow growth or under stress. Many genetic studies on *E. coli* and budding yeast used the term “adaptive mutation” (11, 12) as they used non-lethal selective conditions for an essential amino acid, nucleotide or antibiotic. An important notion related to adaptive mutation is that stress conditions may increase mutations and trigger accelerated evolution (13, 14). A more recent notion is that a bacterial subpopulation of phenotypic variants called ‘persisters’ are more resistant to stress conditions suggesting that they precede adaptive mutations (for review (15)), a notion that was also found in budding yeast (13). The survival of ‘persisters’ to a large range of stress conditions can be explained by a reduced growth rate and metabolism. In addition, life alternates between periods of cell division and quiescence. During quiescence, the main replication-dependent source of mutations is not applicable but others remain, such as DNA repair errors, that may potentially improve the chance of survival. In other words, the respective fitness of cell division and quiescence might alternatively subject organisms to natural selection. In this context, the extent of the impact of replication-independent mutations on the overall mutation load and evolution is unknown. Hence, because of its mechanistic difference, a replication-independent mutational spectrum is expected to exhibit a different signature.

Quiescence is a common cell state on earth (16). In metazoan, stem cells alternate between variable periods of growth and quiescence depending on the period of development and the type of tissues. For germ cells, the male gametes explore the mitotic potential whereas the female are spending long periods in quiescence. For the unicellular fission yeast, *Schizosaccharomyces pombe*, removing nitrogen triggers mating of opposite mating-types followed by meiosis. However, when the population is composed of only one mating-type they arrest in the G1-phase and rapidly enter into quiescence with a 1C content (17). In these conditions, the cells remain viable for months given the medium is refreshed every other week. They are metabolically active, exhibit stress-responsive signaling and are highly efficient in DNA damage repair (18-22). Thus, quiescence in fission yeast is defined under a simple nutritional change so that studies can be reproduced and interpreted rigorously.

Here, we report the accumulation and spectrum of spontaneous mutations that arise in the quiescent phase of fission yeast. The growth and quiescence mutational spectra exhibit quantitative and qualitative differences, that further explore the genetic potential of the genome. We named the new quiescence mutational spectrum “Chronos” the personification of time in Greek mythology.

## Results

In all our experiments, a prototrophic progenitor fission yeast strain is grown in minimal medium (MM) (23) prior to transfer into MM lacking nitrogen at a cell density of 10^6^ cells per milliliter. After two divisions, a majority of the cells arrests in the G1 phase and enters into the G0 quiescent state. The efficiency and accuracy of the repair of DNA lesions in quiescence remain unknown, and lesions are converted into mutations either during quiescence or when cells re-enter the vegetative cycle.

We determined the mutation frequencies by plating samples of quiescent cultures after 1, 4, 8, 11 or 15 days in MM lacking nitrogen onto rich medium containing 5-fluoroorotic acid (5-FOA) that allows the recovery of *ura4^-^* and *ura5^-^* loss-of-function mutants (24). We ascertained that the *ura4*Δ mutants remain viable for two weeks of quiescence, indicating that colonies resistant to 5-FOA (FOA^R^) emerging early are not biased by selection (Figure 1-figure supplement 1) and that our phenotypic accumulation assay is unbiased during the course of the experiment. FOA^R^ colonies were scored and their DNA isolated for mutational spectrum analysis by Sanger sequencing. At day 1, a large fraction of FOA^R^ colonies derive from replicative mutations that have appeared during the last rounds of DNA replication prior or during entry into quiescence and are capable of surviving two to three generations on media lacking uracil. Since such mutations can be found multiple times, mutations found more than once in clonal population were discarded (Figure 1-table supplement). Thus, the frequency of non-redundant FOA^R^ mutations at day 1 ranges from 1 to 3 × 10^−7^ across the various independent experiments (Figure 1A). We next analyzed their spectrum at day 1 by sanger sequencing the respective *ura4^-^* or *ura5^-^* mutated gene. If every substitution occurs with an equal probability, we should observe one transition per two transversions. We found a 1:2.5 ratio, with a mutational bias towards the enrichment of A/T (1.84, Table 1) due to the high frequency of C:G to T:A transitions and G:C to T:A transversions (Figure 1B). Interestingly, as previously observed (5-7) the CpG dinucleotides are found more mutated than other dinucleotides (Figure 1-figure supplement 2) a feature difficult to understand in the absence of cytosine methyl transferase in fission yeast. Among the *ura4^-^* and *ura5^-^* FOA^R^ mutations, 72% are caused by Single Nucleotide Variants (SNVs) and 28% by insertions/deletions (indels) (Figure 1C). A slight bias for insertions is observed with a net gain of 269 bp for 24 events and loss of 334 bp for 16 events (Table 1), including two deletions of 165 and 95 base-pairs (Supplementary Excel file 1). Overall, the mutation profile at day 1 is similar to published results in cycling cells for *URA3* in budding yeast (25) and for *ura4*^+^ and *ura5*^+^ in fission yeast (26).

**Table 1:** Type of mutation and AT bias found in *ura4* and *ura5* mutants at various time points of quiescence

|  | day 1 | day 8 | day 15 | day 8+15 |
| --- | --- | --- | --- | --- |
| **# of SNVs** | **105** | **107** | **37** | **144** |
| Ts | 33 | 26 | 5 | 30 |
| Tv | 72 | 81 | 34 | 116 |
| Ts:Tv | 1:2.18 | 1:3.12 | 1:6.67 | 1:3.70 |
| To A or to T | 63 | 45 | 15 | 60 |
| To C or to G | 17 | 30 | 8 | 38 |
| AT bias | 3.71 | 1.50 | 1.88 | 1.58 |
| **# of indels** | **40** | **139** | **86** | **220** |
| # of deletions | 16 | 99 | 71 | 170 |
| lost bases | 334 | 864 | 907 | 1771 |
| # of insertions | 24 | 35 | 15 | 50 |
| gained bases | 269 | 401 | 99 | 500 |
| net loss | 65 | 463 | 808 | 1271 |
| # of deletions / # of insertions | 0.7 | 2.8 | 4.7 | 3.4 |
| average loss per event | 21 | 9 | 13 | 10 |
| average gain per event | 11 | 11 | 7 | 10 |

Ts: transitions; Tv: transversions AT bias: (GC→AT + GC→TA) to (AT→GC + AT→CG) mutations

**Figure 1.**
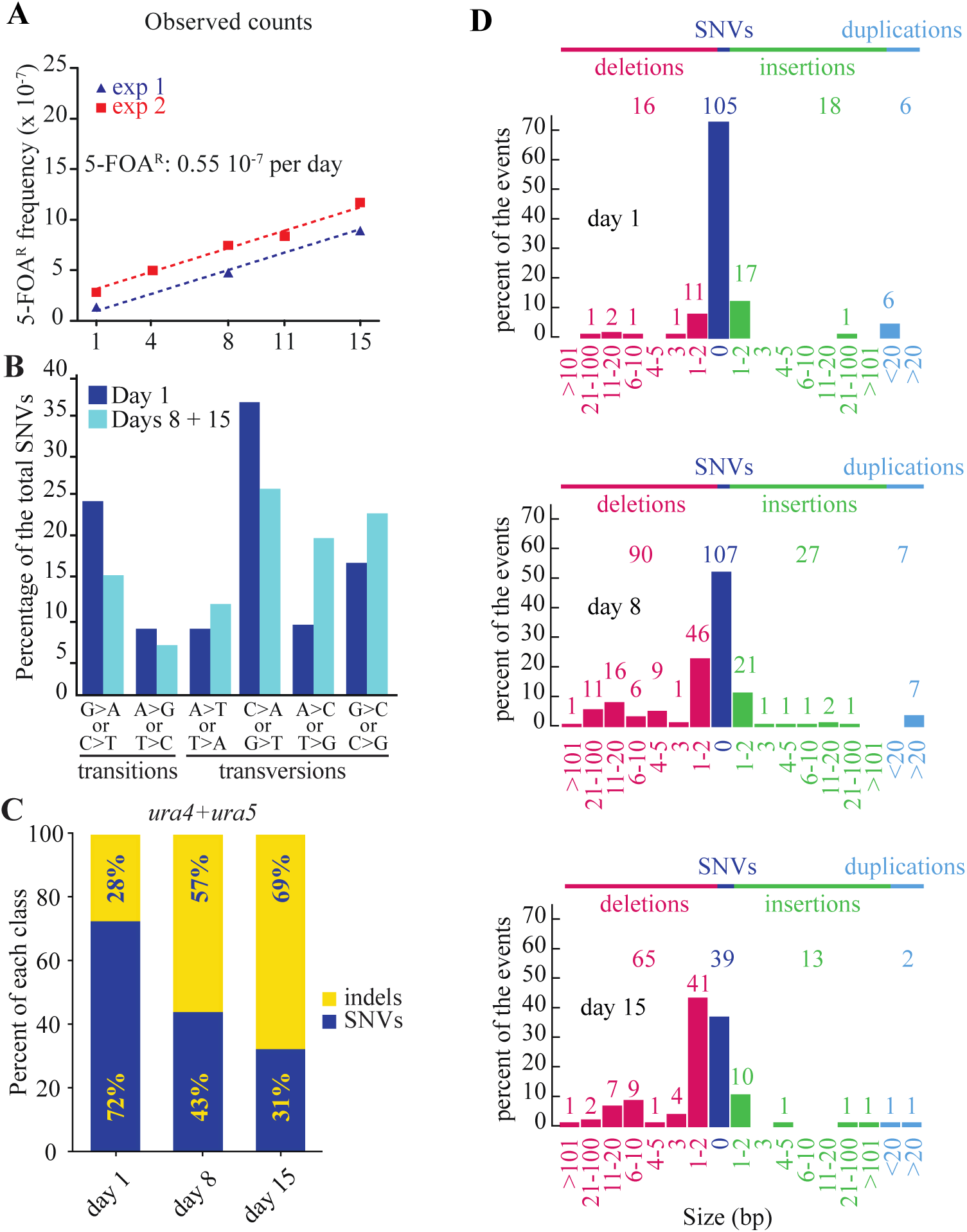
Mutations accumulate as a function of time in quiescence. (**A**) slopes of FOA^R^ accumulation as a function of time in quiescence from two independent experiments, determined by the least squares regression. (**B**) mutation spectrum established on 105 SNVs found in the *ura4*^+^ and *ura5*^+^ genes at day 1 and 146 at days 8 + 15. (**C**) distribution among indels and SNVs of the mutations that result in FOA^R^ for *ura4*^+^ and *ura5*^+^ genes over time. (**D**) distribution of the various sizes of indels over time in quiescence. The numbers directly above the histograms indicate the number of events, while those on the top of the figure are the sum of all the events for each class. The numbers at the bottom indicate the size range of the indels. Figure 1-table supplement shows the position and nature of the substitutions and indels yielding 5-FOA resistant colonies at day 1, 8 and 15. Figure 1-figure supplement 1 shows the survival of wild-type, *ura4D18* cells and a 50% mixture of both cultures were put into quiescence and their survival was followed for two weeks. Figure 1-figure supplement 2 shows (**A**) Percentage of each dinucleotide in the open reading frames of *ura4*^+^ and *ura5*^+^. (**B**) Percentage of each dinucleotide in the combined open reading frames of *ura4*^+^ and *ura5*^+^. (**C**) Observed to expected ratio for each dinucleotide in the open reading frames of *ura4^-^* and *ura5^-^* FOAR clones. (**D**) Observed to expected ratio for each dinucleotide in the combined open reading frames of *ura4^-^* and *ura5^-^* FOAR clones.

During quiescence, the number of mutations resulting in FOA^R^ colonies increases linearly as a function of time. From multiple experiments, we used least squares regression to determine that the slope is 0.55 × 10^−7^ FOA^R^ mutants per day spent in quiescence (Figure 1A). Importantly, we observed that the number of redundant mutations dramatically fades over time, indicating that novel mutations arise in quiescence (Figure 1-table supplement). By combining the SNVs arising after 8 and 15 days of quiescence, the ratio of transition-to-transversion was reduced from 1:2.5 to 1:3.8 and the mutational bias toward A/T decreased (1.84 vs 1.09, Table 1). This is mainly due to a relative decrease of the G:C to A:T transitions that is balanced by an increase of the A:T to C:G and G:C to C:G transversions (Figure 1B). During two weeks of quiescence, the proportion of *de novo* indels increases from 28% at day 1 to 57% and 69% after 8 and 15 days, respectively (Figure 1C) to outnumber the SNVs (Figure 1C and 1D). Altogether, we found more deletions than insertions with a net loss of 1771 bp in 170 mutants and a gain of 500 bp in 50 mutants (Figure 1D, Table 1). The main class of indels is ±1 bp and accounts for roughly half the events (Fig. 1D). More than one half of the indels occur within and near low complexity sequences and several mutations are complex (Figure 1-table supplemental). Collectively, we found a phenotypic mutation rate of 0.55 × 10^−7^ FOA^R^ colonies per day of quiescence with 0.14 × 10^−7^ FOA^R^ colonies per day due to SNVs and 0.41 × 10^−7^ FOA^R^ colonies per day due to indels (Figure 2A and 2B).

**Figure 2.**
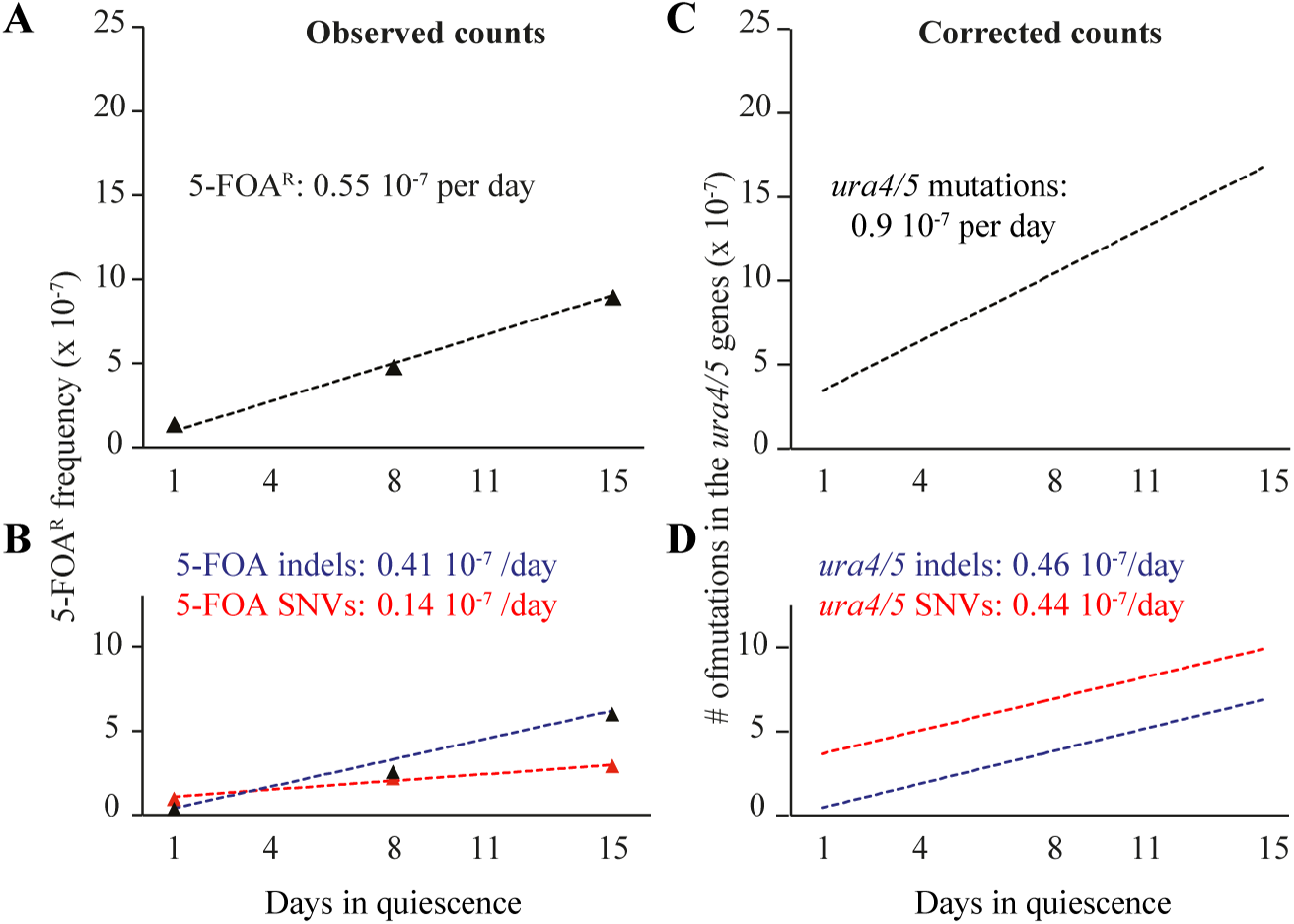
FOA^R^ accumulation and corrected counts. (**A**) slope of FOA^R^ (as determined in Figure 1A). (**B**) slopes of FOA^R^ SNVs and indels accumulation. (C) corrected counts determined on all experimental points using least squares regression (D) corrected slopes for SNVs and indels formation in the *ura4*^+^ and *ura5*^+^ genes. Figure 2-equations supplement shows the methods used to estimate the fraction of essential amino-acids in ura4 and ura5. Figure 2-table supplement 1 shows the estimation of the fraction of SNVs and deletions that exhibit a phenotype. Figure2-table supplement 2 shows the estimation of the fraction non-synonymous SNVs that exhibit a phenotype.

Thus, the mutational spectra of growth and quiescence exhibit striking quantitative and qualitative differences. First, elevated levels of indels are generated. Second, two recent MA studies (5, 6) with fission yeast have shown that in cycling cells insertions outnumber deletions, whereas we observe the reverse in quiescence (Figure 1D). Thus, growth and quiescence apply opposite pressures on the *S. pombe* genome size. Third, during cell divisions, SNVs elevate the genomic A/T composition (8, 9). This bias is reduced during quiescence and counteracts the universal A/T enrichment observed during cell division. This suggests that, in addition to the previously hypothesized gene conversion, nucleotide modifications or transposition, the equilibrium of size and composition of the *S. pombe* genome also depends on the relative strength of the opposing forces applied during growth and quiescence.

Several methods inferring the total mutation count from the measured 5-FOA^R^ phenotypic mutation rate have been described, both for SNVs and indels (11, 25). Here we developed a novel method that relies on the estimation of the fraction of essential amino-acids from the distribution of the number of independent phenotypic mutations observed per amino-acid (Figure 2-equations supplement). When applied to the *ura4*^+^*/5*^+^ genes, we found that the corrected number of mutations per day is 0.9 × 10^−7^, taking into account the mutations that do not result in a 5-FOA^R^ phenotype. When applied to SNVs and indels, we found very similar slopes of 0.445 × 10^−7^ and 0.459 × 10^−7^, respectively (Figure 2D, table supplement 1). Next, we used this method to extrapolate the number of mutations in the quiescent genome. We expected to find 0.904 × 10^−7^ / 1400 (nucleotides of *ura4* + *ura5*) × 14 × 10^6^ (nucleotides in the *S. pombe* genome) = 0.904 × 10^−3^ mutations per genome per day.

To extend our observations made on FOA^R^ mutations to the whole genomes, we analyzed the mutation spectrum in cells that survived for 3 months of quiescence. For long-term experiments, we changed the medium every other week to maintain oligo elements and glucose as well as to prevent the survivors to feed on the nitrogen released by dead cells. In these conditions, the viability at three months is about 0.05% (Fig. S3). From several experiments, we observed a biphasic viability curve, with a cell death acceleration after three weeks of quiescence (Fig. S3). DNA from 123 colonies was purified. Illumina libraries were constructed and paired-end sequenced with an average coverage above 60x to maintain high quality sequences and a low false discovery rate (FDR) that was experimentally validated (see Materials and Methods). SNVs and short indels were determined using Genome Analysis Toolkit (GATK) (27-29), and we combined the output of several tools including SOAPindel, Prism and Pindel (30-32) to increase the sensitivity of indels detection. We performed a stringent calling procedure for both SNVs and indels and we only considered variants that are present in at least 40% of the reads with a local coverage above 10x. Sanger sequencing was used to validate the *de novo* variants and to estimate the FDR.

We report 80 unique mutations, including 40 SNVs and 40 indels from the 123 sequenced genomes (Table 2-table supplement 1). Although low, the mutation level after 3 months was 7 times higher than anticipated by our projection, indicating that this process is not linear for extended periods of time, as suggested by the viability curve (Fig. S3). Among the 29 SNVs involved in the AT bias, we found an AT/GC ratio bias of 1.42, a value intermediates between 1.84 at day 1 or 1.1 at days 8 + 15. This value is higher than those observed in our targeted experiments, but lower than in cycling cells. Among the 40 indels, all the events detected with GATK were found as well with either Pindel, SOAPindel or Prism (30-32). Among these indels, 17 are deletions (17/40 – 42%), a value >2 times greater than in cycling cells (1/6 – 17%) (5).

**Table 2:**
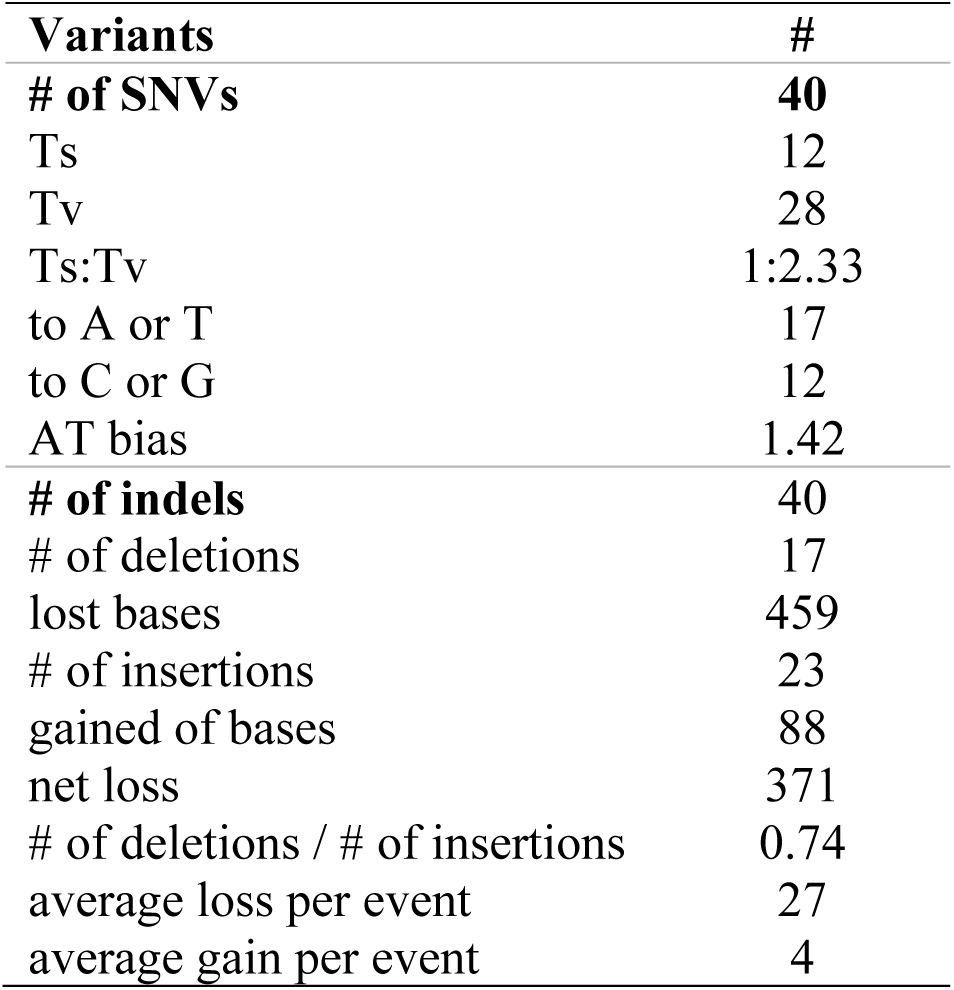
Type of mutation in whole genomes after 3 months in quiescence

Table 2-figure supplement 1 shows the survival curves (red: linear; blue: log) of prototrophic cells in G0 as a function of time. The medium is being replaced every other week starting at day 15, while an aliquot is plated out to monitor the viability. The standard error of the experiments is indicated.

Table 2-table supplement 1 shows the position and nature of the substitution and indels in the 123 genomes.

Table 2-source data. The data has been registered with the BioProject database.

SubmissionID: SUB2493387

BioProject ID: PRJNA379335

http://www.ncbi.nlm.nih.gov/bioproject/379335

Consistent with what was observed in the FOA^R^ study, over 67% (27/40) of the insertions and deletions are ± 1 nucleotide. With respect to genome size, the 40 indels led to the net loss of 371 (459-88) nucleotides. Therefore, the long-term quiescence experiment results support (i) the comparable amounts of indels and SNVs predicted in our estimation (ii) that the proportion of deletions among indels is higher than in cycling cells, with more nucleotides lost than gained (iii) over 67% of the deletions/insertions are ±1 events.

The level of mutations that we have observed after three months in quiescence is likely to cause some heritable phenotypic diversity. Therefore, we conducted a phenotypic survey in conditions that affect a broad range of cellular functions. We did not observe any phenotype upon examination of 384 colonies after 1 day or 1 month of quiescence. However, after 2 and 3 months, we observed 4/376 (1.1%) and 6/334 (1.8%) colonies displaying phenotypes (Table 3), respectively. Genetic crosses confirmed that the phenotype observed in the 10 colonies derives from a single mutated locus. Taken together, we conclude that cellular quiescence allows for genetic variation.

**Table 3:** Phenotypic alterations as a function of time spent in quiescence

| Time<br>in G0 | # of colonies<br>w/phenotype | 18°C | 37°C | KCl<br>1M | Ca(NO<sub>3</sub>)<sub>2</sub><br>0.15M | CaCl<sub>2</sub><br>0.3M | TBZ<br>15µg/ml | HU<br>4mM | SDS<br>0.01% |
| --- | --- | --- | --- | --- | --- | --- | --- | --- | --- |
| 1 day | 0/384 | - | - | - | - | - | - | - | - |
| 1 month | 0/384 | - | - | - | - | - | - | - | - |
| 2 months | 4/376 | - | - | 1 | 1 | 4 | - | - | - |
| 3 months | 6/334 | 1 | 1 | 1 | 1 | 3 | 3 | 2 | 1 |

(-) indicates the absence of a selected phenotype (all the cells form colonies) The numbers in the columns indicate the number of colonies exhibiting a sensitivity to a given treatment (some mutants display a sensitivity to multiple treatments). Serial dilutions were spotted on rich medium plates containing (or not) the drugs at the indicated concentrations. (TBZ): Thiabendazole; (HU): Hydroxyurea; (SDS): Sodium Dodecyl Sulfate

## Discussion

We are aware that the analysis of the 123 genomes analyzed after 3 months of quiescence may not provide the strongest statistical significance. Nonetheless, our results indicate that the *S. pombe* genome fluctuates between two mutational spectra, which alternatively expose the genome to natural selection that progressively shapes its composition, size and ability. During growth, the universal substitution bias toward AT preference along with the domination of insertions over deletion have been reported in numerous studies (8, 9). The fact that the replication-driven mutational bias has not yet reached an equilibrium strongly suggests the existence of forces capable of counterbalancing it. Here we propose that quiescence is a novel mutational force that together with DNA replication-dependent mutations, DNA repair errors, nucleotide modifications, recombination or transposition impact on the genetic material. Separate and alternate modes of mutagenesis and selection allow compensatory mutations to arise in one phase of the life cycle and fuel the phenotypic evolution simultaneously into the subsequent phase. On the contrary, the super-housekeeping genes (19) constrain evolution with a strong vector of conservation, since they are required for proliferation, quiescence and the transition from one to the other (33).

Concerning the role of mutations in evolution and the central position of the gametes, taking into account that fission yeast is haploid and sexually competent, it is tempting to propose that in multicellular organisms each gamete specifically selects for high efficiency of either proliferation or quiescence. These qualities eventually merge in the diploid zygote, prior to the initiation of a new life cycle, suggesting that the fundamental difference in the lifestyle of the two gametes contributes to the genetic fitness of the dividing and quiescent somatic cells and the homeostasis of adult tissues with aging. In return, it may participate to the dichotomy in gamete type and size found in multicellular organisms. A consequence of our model is that the spectrum of variants observed on the sexual chromosomes of mammals differ from that of the autosomes (Achaz et al. submitted). To our knowledge, James Crow (34) was the first to hypothesize a higher occurrence of indels, in particular deletions, in the female gamete lineage than in her male counterpart. In this regard, we suspect that the origin of mutations is paternally biased mostly because SNVs were predominantly taken into account. However, the recent progress in indels detection with new generation technologies will undoubtedly reveal many overlooked indels (35).

The genetics of quiescence underscores the importance of a replication-independent but time-dependent process (36-39), to the overall mutation spectrum. This time-dependent process should also help to fine-tune the accuracy of the “molecular clock” that measures, in units of time (40), the evolutionary distance of two closely related species with their common ancestor. Such hypotheses are accessible to experimental and modeling approaches and are of great interest for evolutionary, developmental and human-health perspectives.

## Materials and Methods

### Strain and Sanger Sequencing of ura4 and ura5 Mutants

The stable prototrophic M-smt0 PB1623 strain was used in all our experiments. We identified the FOA^R^ mutant strains as *ura4^-^* and *ura5^-^* mutants by genetic crosses with a known *ura4*Δ strain, purified their DNA, PCR amplified the gene of interest, analyzed the PCR fragment by agarose gel electrophoresis and subjected it to Sanger sequencing.

### Library Construction and Sequencing

Library construction and sequencing was performed by Illumina HiSeq 2500 following the manufacturer’s instruction. Base calling was performed using CASAVA 1.9. For each strain, one insert size (ranging from 400 to 800 bp) library was constructed and sequenced. After initial quality control assessment with FastQC version 0.10.1 (41) fqCleaner (l = 80; q = 30) was used to trim the tails of the reads if the Phred quality dropped below 30.

### Alignment-Based Assembly

We sequenced 12 strains per lane on an Illumina HiSeq 2500, aligned the resulting reads to the *Schizosaccharomyces_*pombe.ASM294v2.23 DNA reference genome with BWA-MEM; version 0.7.5a. (42). SAMtools version 0.1.19 and Picard version 1.96 (http://picard.sourceforge.net) were used to process the alignment files and to mark duplicate reads. The coverage in our experiments ranges from 60 to 120, with an average of 80. SNVs and small indels were called using GATK version 2.7-2 (27). We applied quality score recalibration, indel realignment, duplicate removal, and performed SNV and INDEL discovery and genotyping using standard filtering parameters or variant quality score recalibration according to GATK Best Practices recommendations (28, 29). In addition to the GATK analysis, we combined three programs, Pindel (32), Prism (31), and SOAPindel (30) dedicated to the detection of INDELs to search for additional variants not detected by GATK. Only variants detected at least 10-times in a sample and not found in any other strain sequenced from the same G0 pool were considered.

### Filtering

For GATK, Pindel, Prism and SOAPindel analyses, only variants detected at least 10-times in a sample and not found in any other strain sequenced from the same G0 pool were considered. For GATK and indels detection, we have determined the FDR by Sanger sequencing on a larger set of previously sequenced strains. 20 random SNVs whose quality scores ranged from 21 to 1700 were analyzed. Only the lowest score (21; one occurrence) turned out to be a false positive, yielding an FDR of 0.05. Concerning the indels, all the variants detected by at least two approaches, including GATK, turned out to be true by Sanger sequencing. For the variants only detected by Prism, we sequenced 13 occurrences and found that only the two lowest scores (DP10) were false positives (FDR = 0.15). Pindel yielded the poorest yield of variants that were systematically true and called by at least one other program. SOAPindel called a large number of variants that were dispatched into 5 classes according to their type of output in the VCF file. We kept only the relevant calls labeled as HP=A_B or HP=X_N in the vcf file for which the FDR determined on 14 variants was 0.2143 (in the three other classes, 54 occurrences, only five were true and were not taken into account).

### Miscellaneous

Template DNA fragments were hybridized to the surface of paired-end (PE) flow cells (HiSeq 2500 sequencing instruments) and amplified to form clusters using the Illumina cBotTM. Paired-end libraries were sequenced using 2 × 120 cycles of incorporation and imaging with Illumina SBS kits. For the HiSeq 2500, 2×101 cycles with SBS kits v3 were employed. Each library was initially run, assessing optimal cluster densities, insert size, duplication rates and comparison to chip genotyping data. Following validation, the desired sequencing depth (> 60X) was then obtained. Real-time analysis involved conversion of image data to base-calling in real-time.

## Figure 1-table supplemental

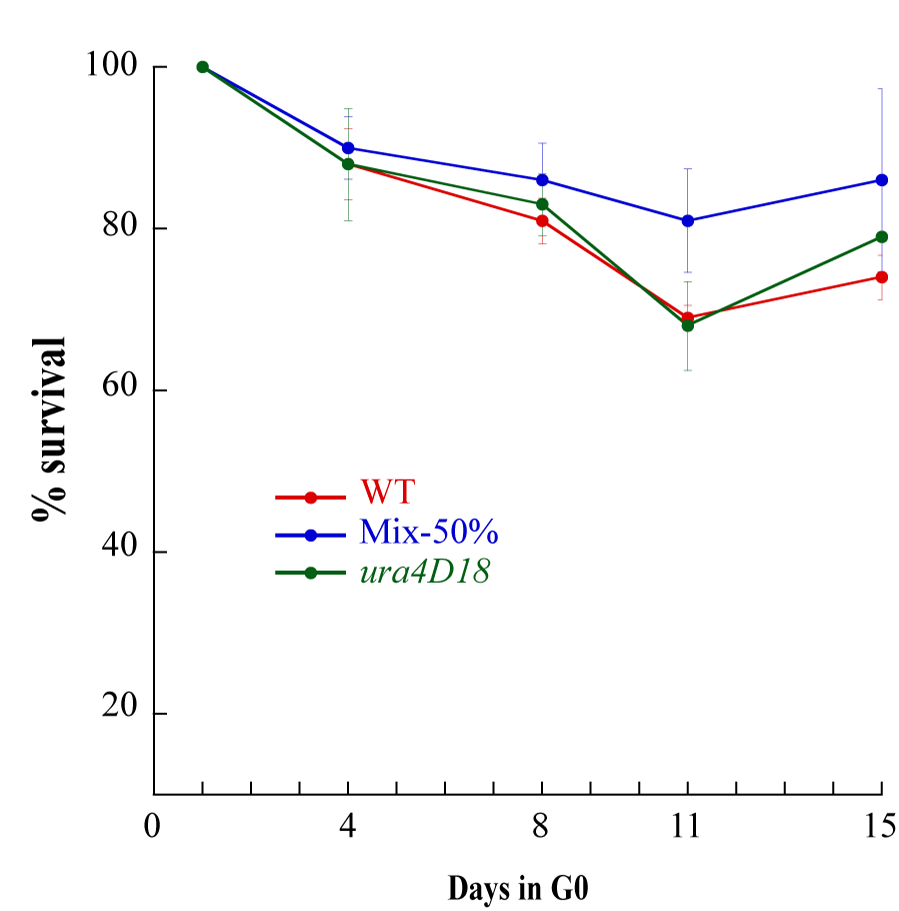
Figure 1-figure supplementary 1. Figure 1-figure supplementary 1 shows the survival of Wild-Type, *ura4D18* cells and a 50% mixture of both cultures were put into quiescence and their survival was followed for two weeks.

### Figure 1-figure supplementary 2

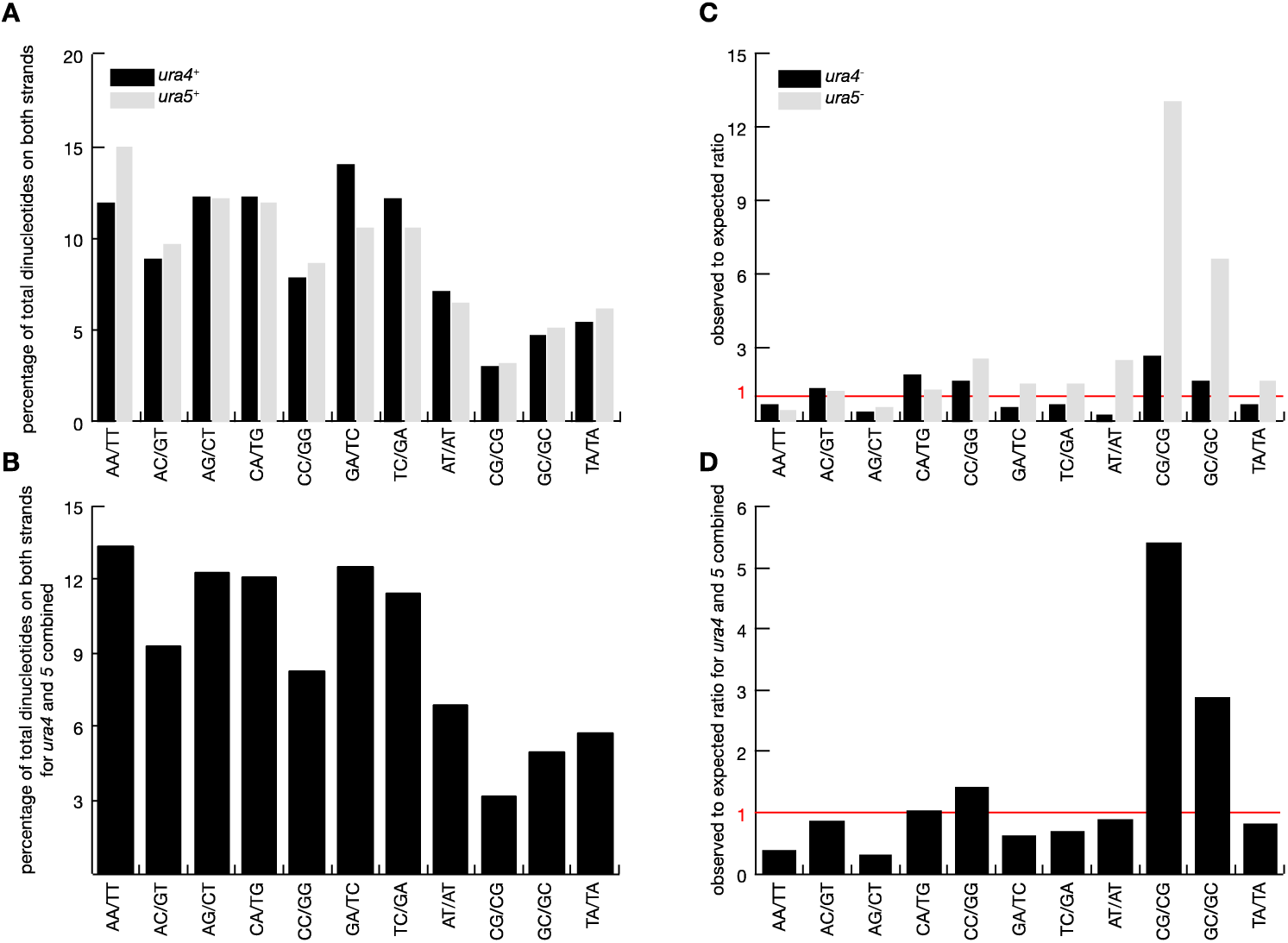
Figure 1-figure supplementary 2. (**A**)Percentage of each dinucleotide in the open reading frames of *ura4*^+^ and *ura5*^+^ (**B**) Percentage of each dinucleotide in the combined open reading frames of *ura4*^+^ and *ura5*^+^ (**C**) Observed to expected ratio for each dinucleotide in the open reading frames of *ura4^-^* and *ura5^-^* FOAR clones (**D**) Observed to expected ratio for each dinucleotide in the combined open reading frames of *ura4*^-^ and *ura5*^-^ FOAR clones.

## Figure 2-equations supplement

### Estimation of the Fraction of Essential Amino-Acids

The experimental assay based on 5-FOA resistance can be used to estimate the rate of mutations that give rise to a phenotype. All the mutations that we have observed are single mutation events that invalidate either the *ura4*^+^ or *ura5*^+^ genes, hence resulting in the FOA^R^ phenotype. We now want to infer the total mutation rate, both for SNVs and indels independently.

We hypothesize that most indels will be deleterious to the genes and that therefore the global indel rate is close the indel rate that results in FOA^R^ (for genes - 50% of the genome). On the contrary, many SNVs are likely to exhibit no phenotype and, therefore, we observe only a fraction of the total number of SNVs.

We have inferred the global SNV rate from the phenotypic mutation rate using two independent methods. Substitutions can be classified as synonymous (i.e. when the amino-acid sequence is unchanged), non-synonymous and STOP mutations. Both methods assume that all STOP mutations yield a phenotype, that all synonymous mutations don’t and aim at estimating 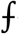_NS_, the fraction of non-synonymous mutations that gives rise to a phenotype. The first one, described by Lang and Murray (25), makes the assumption 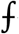_NS_ can be estimated directly from the fraction of observed STOP mutations in the sample to the total number of potential STOP in the sequence. The second method, developed for our purpose, assumes that the amino-acids can all be classified as essential (once mutated the function is lost) or not and that 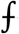_NS_ can be assimilated to the fraction of essential amino-acids in the sequences. We estimated the fraction of essential amino-acids from the distribution of the number of independent FOA^R^ mutations observed per amino acid (hereafter the *k*-distribution). The results of both methods are reported in Table S1. We used the difference between the predicted and observed *k*-distribution as a goodness of fit. We find that both methods yield similar results but that our method outperforms the one proposed by Lang and Murray (25). Hence, we only used our estimation in the course of the study.

A gene can be modeled as a sequence of *K* amino acids, among which only *k* are essential. If *M* non-synonymous mutations occur on this gene, only a subset *m* will target the *k* essential amino acids. If *k* were known, we could estimate *M* as *mK/k*. Therefore, since *K* and *m* are observed, we need to estimate *k* in order to estimate *M*. In the following, we estimated *k* by maximum likelihood using the distribution of observed point mutations per essential amino acid (i.e. the k-distribution). We observe a total of *m* mutations that are distributed among the *k* essential amino acids, where amino-acid *i* has *m*_i_ mutations:

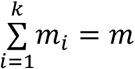

As each mutation has a probability 1/*k* to occur at a particular essential amino acid, the set of *m*_i_ is given by the multinomial probability distribution:

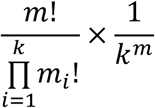

However, we consider in our model that amino acids with the same number of substitutions are exchangeable. Defining *k_j_*, as the number of amino acids that have *j* substitutions, the number of exchangeable configurations is given by:

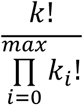
where *max* is the highest observed number of substitutions for an amino acid. Therefore, once rearranged, the likelihood of the overall observed distribution of (*k*_1;_ *k*_2_,… *k*_max_) given *k* is:

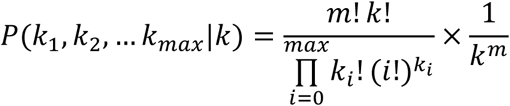
with

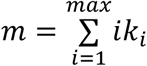
and

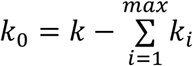

We thus computed numerically the maximum likelihood estimate as well as its associated 95% credibility interval. All counts as well as 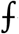_NS_ estimates using both methods are reported in Table S1. The goodness-of-fit tested the difference between the k-distribution predicted from 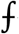_NS_ to the observed k-distribution using a chi-2 test. The expected frequencies in the -distribution is given by a binomial (*1/k*, #non-synonymous substitutions) conditioned to be non-zero, as essential amino acids that were not mutated cannot be observed. This test shows that the fractions estimated by the Lang and Murray method cannot explain the observed distribution. Since we observe only substitutions that disrupt the function of the genes, we assume that all synonymous substitutions have no phenotype but that all STOP substitutions do as well as a fraction 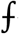_NS_ of the non-synonymous substitutions.

We computed the distribution of deletion size and corrected by 1/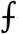_NS_ deletions of size 3 bp -the deleted amino acid is essential– and 1/ (1-(1-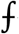_NS_)^2^) for the deletions of size 6 bp –at least one of the two deleted amino-acid is essential–. We assume that all other deletions lead to a loss of function and that insertions follow the same logic. Results for both estimates are reported in Table S2.

## Figure 2-table supplement 1

**Table S1:** Estimation of the fraction of non-synonymous substitutions that exhibit a phenotype.

|  |  | *ura4*<br>(265 Aa) | *ura5*<br>(216 Aa) | total<br>(481 Aa) |
| --- | --- | --- | --- | --- |
| Lang & Murray<br>2008 | Fraction of substitutions<br>with a phenotype | 0.22 | 0.18 | 0.20 |
|  | Fraction of deletions<br>with a phenotype | 0.73 | 0.96 | 0.85 |
| ML method | Fraction of substitutions<br>with a phenotype | 0.39 | 0.23 | 0.30 |
|  | [CI 95%] | [0.30,0.61] | [0.19,0.32] | [0.26,0.39] |
|  | Fraction of deletions<br>with a phenotype | 0.83 | 0.98 | 0.90 |

## Figure 2-table supplement 2

**Table S2.**
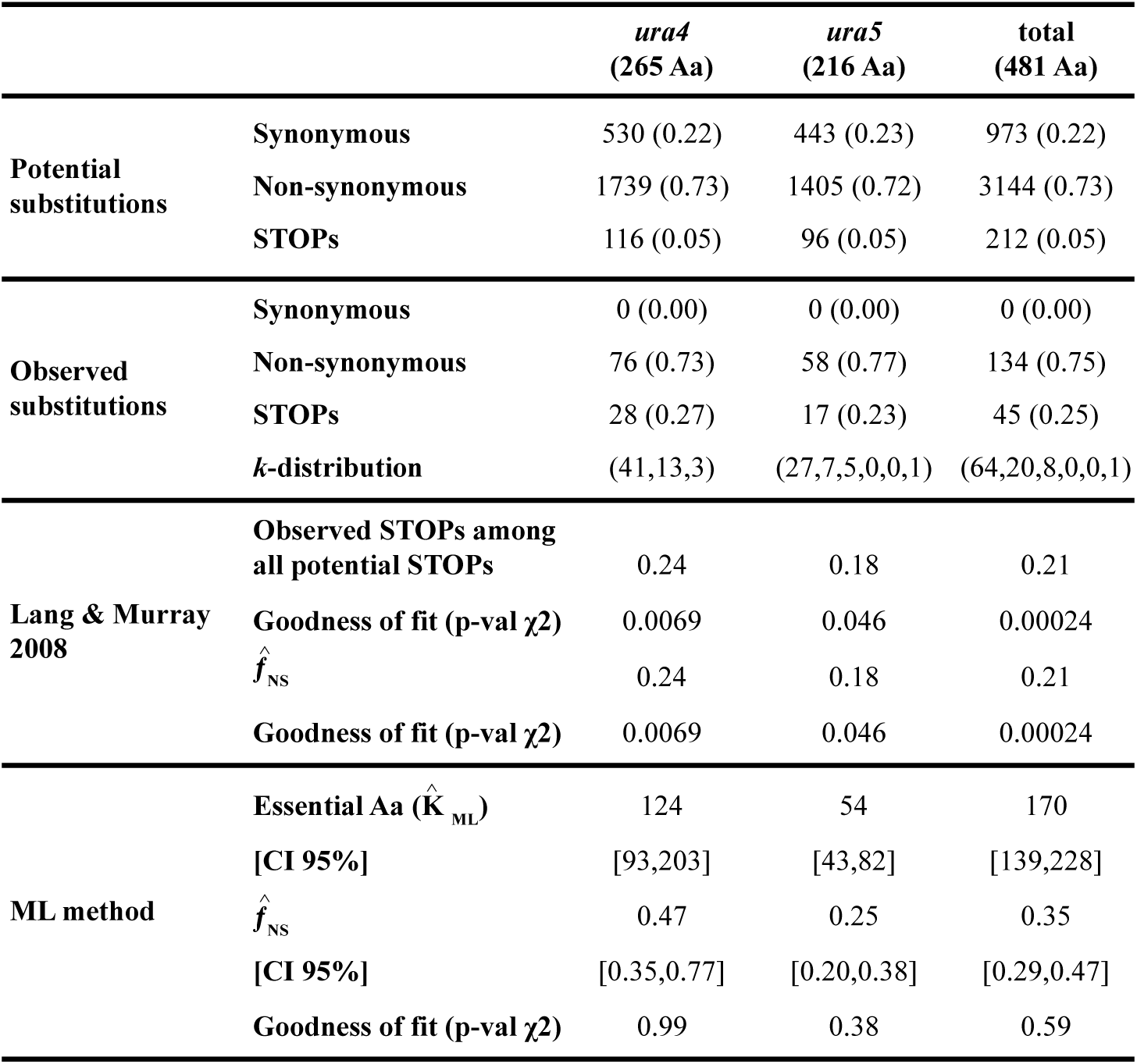
Estimation of the fraction of non-synonymous substitutions that exhibit a phenotype

|  |  | *ura4*<br>(265 Aa) | *ura5*<br>(216 Aa) | total<br>(481 Aa) |
| --- | --- | --- | --- | --- |
| Potential<br>substitutions | Synonymous | 530 (0.22) | 443 (0.23) | 973 (0.22) |
|  | Non-synonymous | 1739 (0.73) | 1405 (0.72) | 3144 (0.73) |
|  | STOPs | 116 (0.05) | 96 (0.05) | 212 (0.05) |
| Observed<br>substitutions | Synonymous | 0 (0.00) | 0 (0.00) | 0 (0.00) |
|  | Non-synonymous | 76 (0.73) | 58 (0.77) | 134 (0.75) |
|  | STOPs | 28 (0.27) | 17 (0.23) | 45 (0.25) |
|  | *k*-distribution | (41,13,3) | (27,7,5,0,0,1) | (64,20,8,0,0,1) |
| Lang & Murray<br>2008 | Observed STOPs among<br>all potential STOPs | 0.24 | 0.18 | 0.21 |
| | Goodness of fit (p-val $\chi^2$) | 0.0069 | 0.046 | 0.00024 |
| | $\hat{f}_{NS}$ | 0.24 | 0.18 | 0.21 |
| | Goodness of fit (p-val $\chi^2$) | 0.0069 | 0.046 | 0.00024 |
| ML method | Essential Aa ($\hat{K}_{ML}$) | 124 | 54 | 170 |
|  | [CI 95%] | [93,203] | [43,82] | [139,228] |
| | $\hat{f}_{NS}$ | 0.47 | 0.25 | 0.35 |
|  | [CI 95%] | [0.35,0.77] | [0.20,0.38] | [0.29,0.47] |
| | Goodness of fit (p-val $\chi^2$) | 0.99 | 0.38 | 0.59 |

### Table 2-figure supplemental 1

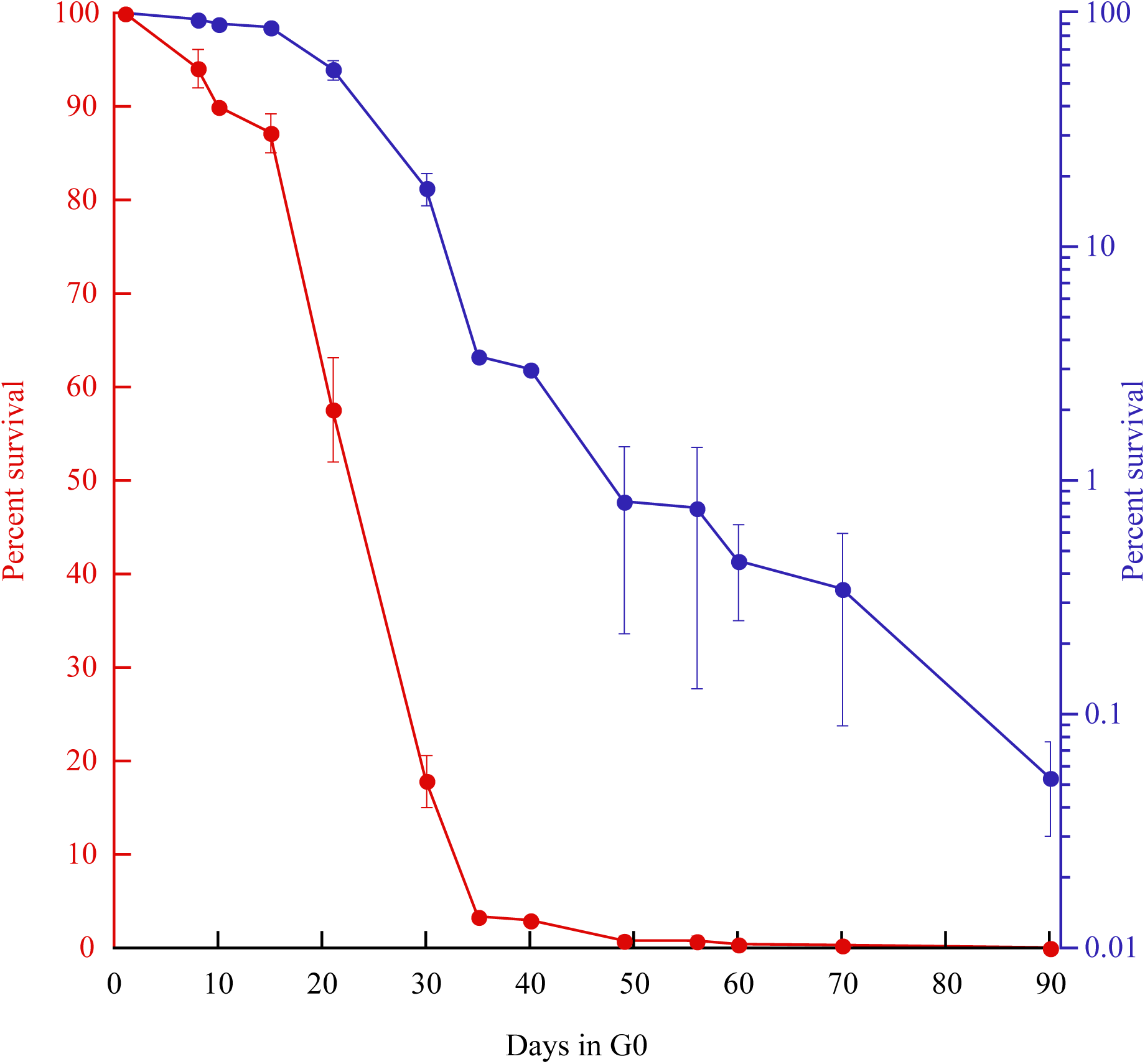
Table 2-figure supplemental 1: Survival curves (red: linear; blue: log) of prototrophic cells in G0 as a function of time. The medium is being replaced every other week starting at day 15, while an aliquot is plated out to monitor the viability. The standard error of the experiments is indicated.

Table 2-table supplement 1 shows the position and nature of the substitution and indels in the 123 genomes. Attached.

Table 2-source data. The data has been registered with the BioProject database.

SubmissionID: SUB2493387

BioProject ID: PRJNA379335

http://www.ncbi.nlm.nih.gov/bioproject/379335

## Acknowledgements

We thank Stefania Francesconi, Karl Ekwall and Stéphane Marcand for their critical reading of the manuscript. This work was supported by the grants ANR-13-BSV8-0018 and ANR-12-BSV7-0012

Demochips from the Agence Nationale de la Recherche (France) to BA and GA, respectively.

BA designed the study. SG and AV performed NGS data analysis. SM and CD performed the bench work. SG, SM and CD performed the Sanger analysis and BA, SG and GA wrote the paper.

## References

1. Luria SE, Delbrück M (1943) Mutations of Bacteria from Virus Sensitivity to Virus Resistance. Genetics 28(6): 491–511.

2. Lea DE, Coulson CA (1949) The distribution of the numbers of mutants in bacterial populations. JGenet 49(3): 264–285.

3. Halligan DL, Keightley PD (2009) Spontaneous Mutation Accumulation Studies in Evolutionary Genetics. Annual Review of Ecology Evolution and Systematics 40(1): 151–172.

4. Conrad DF, et al. (2011) Variation in genome-wide mutation rates within and between human families. Nat Genet 43(7): 712–714.

5. Behringer MG, Hall DW (2015) Genome-Wide Estimates of Mutation Rates and Spectrum in Schizosaccharomyces pombe Indicate CpG Sites are Highly Mutagenic Despite the Absence of DNA Methylation. G3 (Bethesda) 6(1): 149–160.

6. Farlow A, et al. (2015) The Spontaneous Mutation Rate in the Fission Yeast Schizosaccharomyces pombe. Genetics 201(2): 737–744.

7. Zhu YO, Siegal ML, Hall DW, Petrov DA (2014) Precise estimates of mutation rate and spectrum in yeast. Proc Natl Acad Sci USA 111(22): E2310–8.

8. Hershberg R, Petrov DA (2010) Evidence That Mutation Is Universally Biased towards AT in Bacteria. PLoS Genet 6(9). doi:10.1371/journal.pgen.1001115.

9. Lynch M (2010) Rate, molecular spectrum, and consequences of human mutation. Proc Natl Acad Sci USA 107(3): 961–968.

10. Zuckerkandl E, Pauling L (1962) Molecular disease, evolution, and genetic heterogeneity. Horizons in Biochemistry, eds Kasha M, Pullman B (New York), pp 189–225.

11. Drake JW (1991) Spontaneous Mutation. Annu Rev Genet 25(1): 125–146.

12. Foster PL (1999) Mechanisms of stationary phase mutation: a decade of adaptive mutation. Annu Rev Genet 33(1): 57–88.

13. Yaakov G, Lerner D, Bentele K, Steinberger J (2017) Coupling phenotypic persistence to DNA damage increases genetic diversity in severe stress. Nature Ecology &… 1(1): 0016.

14. Long H, Behringer MG, Williams E, Te R, Lynch M (2016) Similar Mutation Rates but Highly Diverse Mutation Spectra in Ascomycete and Basidiomycete Yeasts. Genome Biol Evol 8(12): 3815–3821.

15. Harms A, Maisonneuve E, Gerdes K (2016) Mechanisms of bacterial persistence during stress and antibiotic exposure. Science 354(6318): aaf4268.

16. Lewis DL, Gattie DK (1991) The ecology of quiescent microbes. (ASM American Society for Microbiology News).

17. Nurse P, Bissett Y (1981) Gene required in G1 for commitment to cell cycle and in G2 for control of mitosis in fission yeast. Nature 292(5823): 558–560.

18. Mochida S, Yanagida M (2006) Distinct modes of DNA damage response in S. pombe G0 and vegetative cells. Genes Cells 11(1): 13–27.

19. Yanagida M (2009) Cellular quiescence: are controlling genes conserved? Trends Cell Biol 19(12): 705–715.

20. Ben Hassine S, Arcangioli B (2009) Tdp1 protects against oxidative DNA damage in non-dividing fission yeast. EMBO J 28(6): 632–640.

21. Marguerat S, et al. (2012) Quantitative analysis of fission yeast transcriptomes and proteomes in proliferating and quiescent cells. Cell 151(3): 671–683.

22. Gangloff S, Arcangioli B (2017) DNA Repair and Mutations During Quiescence in Yeast. FEMS Yeast Res. doi:10.1093/femsyr/fox002.

23. Mitchison JM (1970) Physiological and cytological methods for *Schizosaccharomyces pombe*. Methods in Cell Biology. doi:10.1016/S0091-679X(08)61752-5.

24. Grimm C, Kohli J, Murray J, Maundrell K (1988) Genetic-Engineering of Schizosaccharomyces-Pombe - a System for Gene Disruption and Replacement Using the Ura4 Gene as a Selectable Marker. Mol Gen Genet 215(1): 81–86.

25. Lang GI, Murray AW (2008) Estimating the per-base-pair mutation rate in the yeast Saccharomyces cerevisiae. Genetics 178(1): 67–82.

26. Fraser J, Neill E, Davey S (2003) Fission yeast Uve1 and Apn2 function in distinct oxidative damage repair pathways in vivo. DNA Repair (Amst) 2(11): 1253–1267.

27. McKenna A, et al. (2010) The Genome Analysis Toolkit: A MapReduce framework for analyzing next-generation DNA sequencing data. Genome Res 20(9): 1297–1303.

28. DePristo MA, et al. (2011) A framework for variation discovery and genotyping using next-generation DNA sequencing data. Nat Genet 43(5): 491–.

29. Van der Auwera GA, et al. (2013) From FastQ data to high confidence variant calls: the Genome Analysis Toolkit best practices pipeline. Curr Protoc Bioinformatics 43:11.10.1–33.

30. Li S, et al. (2013) SOAPindel: Efficient identification of indels from short paired reads. Genome Res 23(1): 195–200.

31. Jiang Y, Wang Y, Brudno M (2012) PRISM: Pair-read informed split-read mapping for base-pair level detection of insertion, deletion and structural variants. Bioinformatics 28(20): 2576–2583.

32. Ye K, Schulz MH, Long Q, Apweiler R, Ning Z (2009) Pindel: a pattern growth approach to detect break points of large deletions and medium sized insertions from paired-end short reads. Bioinformatics 25(21): 2865–2871.

33. Williams GC, Williams DC (1957) Natural Selection of Individually Harmful Social Adaptations Among Sibs With Special Reference to Social Insects. Evolution 11(1): 32.

34. Crow JF (2000) The origins, patterns and implications of human spontaneous mutation. Nat Rev Genet 1(1): 40–47.

35. Ponting CP, Nellåker C, Meader S (2011) Rapid turnover of functional sequence in human and other genomes. Annu Rev Genomics Hum Genet 12(1): 275–299.

36. Goldmann JM, et al. (2016) Parent-of-origin-specific signatures of de novo mutations. Nat Genet 48(8): 935–939.

37. Hazen JL, et al. (2016) The Complete Genome Sequences, Unique Mutational Spectra, and Developmental Potency of Adult Neurons Revealed by Cloning. Neuron 89(6): 1223–1236.

38. Kumar S, Subramanian S (2002) Mutation rates in mammalian genomes. Proc Natl Acad Sci USA 99(2): 803–808.

39. Ségurel L, Wyman MJ, Przeworski M (2014) Determinants of mutation rate variation in the human germline. Annu Rev Genomics Hum Genet 15(1): 47–70.

40. Reis dos M, Donoghue PCJ, Yang Z (2016) Bayesian molecular clock dating of species divergences in the genomics era. Nat Rev Genet 17(2): 71–80.

41. Chen C-J, et al. (2012) ncPRO-seq: a tool for annotation and profiling of ncRNAs in sRNA-seq data. Bioinformatics 28(23): 3147–3149.

42. Li H, Durbin R (2009) Fast and accurate short read alignment with Burrows-Wheeler transform. Bioinformatics 25(14): 1754–1760.

